# A ubiquitous spectrolaminar motif across independent studies, including Mackey et al.’s own data

**DOI:** 10.1101/2025.05.07.652644

**Authors:** Alex James Major, Ahmed Abdaltawab, Jessica M. Phillips, Tian Wang, Eric Kenji Lee, Maxwell J. Lichtenfeld, Chandramouli Chandrasekaran, Yuri B. Saalmann, Alexander Maier, Robert Desimone, Earl K. Miller, André M. Bastos, Diego Mendoza-Halliday

## Abstract

Our study (Mendoza-Halliday et al., 2024) made two contributions: (1) discovery of a ubiquitous cortical motif and (2) a tool derived from it—the Frequency-based Layer Identification Procedure (FLIP and vFLIP). Mackey et al. critique the tool, questioning its advantage over classic current source density (CSD) analysis, and reason backwards to challenge the motif’s ubiquity. In our rebuttal, we confirm the spectrolaminar motif in diverse cortical areas using data from multiple research groups (who joined us in this rebuttal) as well as Mackey et al.’s own dataset. Additionally, we introduce vFLIP2, an improved version of our tool that addresses their comments. It reliably identified and localized the motif in our data and Mackey et al.’s data. Our findings reaffirm the motif’s ubiquity. We value Mackey et al.’s comments, which helped refine our tool.

## Main Text

Mackey et al. argue that the spectrolaminar motif reported in our study (Mendoza-Halliday et al., 2024) is neither generalizable nor ubiquitous. However, this claim contradicts results from 15 studies across multiple independent labs (Supplementary Table 1). In our original dataset, we observed the motif across hundreds of probe recordings spanning 14 cortical areas (Supplementary Fig. 1)—a sample more than 14 times larger than Mackey et al.’s limited sample of a few dozen probes in just three cortical areas.

Curiously, and despite Mackey et al.’s claims to the contrary, the spectrolaminar motif was present in their own data, which they provided to us. We first analyzed the laminar power profiles of their probe recordings (N = 26 for A1, N = 24 for V1, N = 9 for Belt) using the manual method (Mendoza-Halliday et al., 2024) to identify probes with opposite laminar power gradients in the gamma (75–150 Hz) and alpha/beta (10–19 Hz) ranges. The spectrolaminar motif is obvious in their individual probe recordings (Supplementary Fig. 1, bottom row), including the example shown in Figure 1b of their commentary. It is also evident in the mean spectrolaminar pattern across probes in each cortical area of their data, regardless of whether we used the manual method (Fig. 1a-d, Supplementary Fig. 2a,e) or the current source density (CSD) sink (Supplementary Fig. 2c,g) to identify the middle layer. In the Mackey et al. dataset, the spectrolaminar motif was identified in 65% (17/26) of the probes in A1, 67% (16/24) in V1, and 64% (9/14) in Belt, for a total of 66% (42/64) probes across areas. This is consistent with our original study (61%). These results show that Mackey et al.’s claim that the motif is not present in their data is incorrect. We cannot explain their claim.

**Figure 1.**
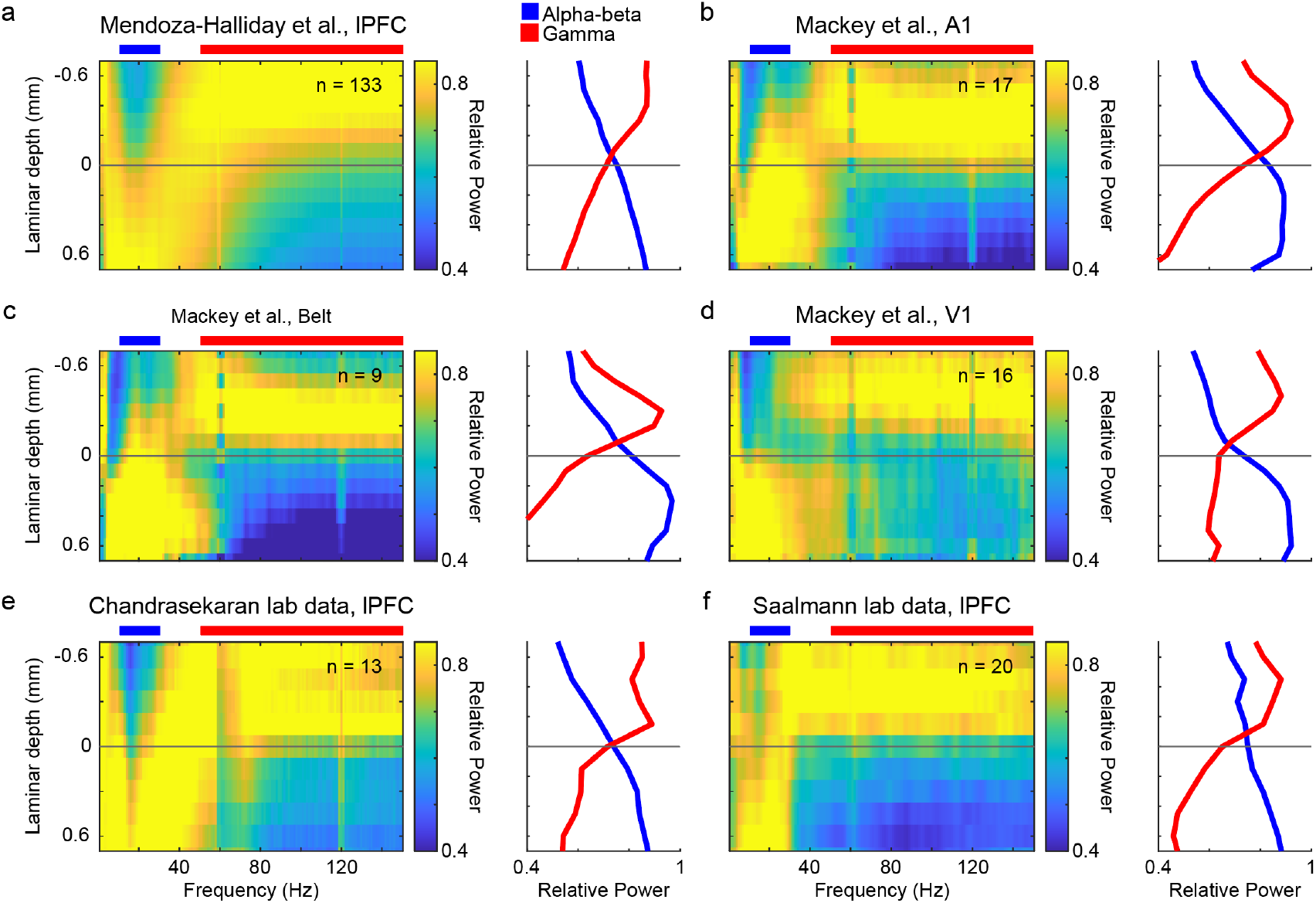
The spectrolaminar motif is ubiquitous across datasets from different labs. For each dataset, across-probes mean relative power map (left) and mean relative power in the alpha-beta (blue) and gamma (red) bands as a function of laminar depth with respect to the crossover channel (right). (a) Dataset from lateral prefrontal cortex (LPFC) probes of original publication (Mendoza-Halliday et al., 2024). (b-d) Data from Mackey et al. collected in primary auditory cortex (A1, b), belt region (c), and primary visual cortex (V1,d). (e) LPFC data provided by Tian Wang and Eric Kenji Lee from the lab of Chandramouli Chandrasekaran, Boston University. f, LPFC data from the lab of Yuri Saalmann (Phillips et al., 2024), University of Wisconsin-Madison. N’s in each spectrolaminar subplot refer to the total number of independent probes that were averaged for each analysis.

One issue may be Mackey et al.’s lack of ground-truthing using anatomical methods. In our study, we used electrolytic lesions across multiple cortical areas to confirm the correspondence between the spectrolaminar motif and histologically identified cortical layers. Mackey et al.’s critique of the anatomical positioning of the spectrolaminar motif relies on the first CSD sink, an indirect electrophysiological proxy for layer 4. This is less accurate than our histologically confirmed measurements. Moreover, their argument that CSD is more accurate, based on CSD itself as the ground truth, is circular. Anatomy remains the only definitive ground truth for a motif rooted in cortical layers.

Further, we confirmed the spectrolaminar motif with data from two other independent research groups, who co-author this rebuttal with us (Fig. 1E, unpublished data from Chandramouli Chandrasekaran, and Fig. 1F, data from Phillips et al., 2024). This new evidence documents a broad consensus on the presence of the spectrolaminar motif in over 20 cortical areas (Supplementary Table 1).

### vFLIP2: A new robust automatic layer identification algorithm

It is puzzling that Mackey et al. question the spectrolaminar motif, as their comments focus primarily on vFLIP—the tool derived from the motif—rather than the motif itself. vFLIP enables real-time identification of cortical layers during experiments and surgeries. While the classic CSD approach remains valuable, vFLIP offers distinct advantages: it does not require strong sensory responses, has a higher signal-to-noise ratio, is more robust to probe angle variations, and is automated. Histology confirmed the improved accuracy in using the spectrolaminar motif for localizing layer 4 (Mendoza-Halliday et al., 2024). Like most tools, it can continue to be refined, and Mackey et al.’s feedback was valuable in developing its next iteration.

We introduce vFLIP2, an enhanced version of the original algorithm. vFLIP2 reliably identifies the spectrolaminar motif and localizes its laminar components in both the Mackey et al. dataset (N = 64 probes) and our original dataset (N = 923 probes). It addresses all the concerns raised by Mackey et al., including risks of false positives from exploring “ a large space of potential frequency bands,” computational inefficiency, the detection of “ very narrow bands” , and incorrect detection of inverted patterns. The algorithm identifies low- and high-frequency ranges with maximal laminar power gradients by computing the relative power difference between proximal and distal channel groups across all frequencies simultaneously, enforcing minimum bandwidth thresholds, and using omega (ω), a more comprehensive metric of gradient quality (see Supplementary Text). This new approach grants vFLIP2 superior computational efficiency, completing spectrolaminar analysis for each probe in an average of 0.7 seconds.

We validated vFLIP2 on both our dataset and the Mackey et al. dataset. Across probes, it consistently detected laminar power gradients in a low-frequency range centered around 16 Hz and a broad high-frequency range in both our data (Figure 2a-c) and Mackey et al.’s data (Supplementary Fig. 3a). The spectrolaminar motif was identified by vFLIP2 in 78% of probes in the Mackey dataset (Fig 2f) and 80% in our dataset (Fig. 2e). The motif is evident in the mean patterns across probes (Fig. 2g; Supplementary Fig. 2b,f). Importantly, vFLIP2 reliably identified the motif in real data but not in layer-shuffled data or pink noise (Fig. 2d,e), demonstrating both sensitivity and specificity. vFLIP2 also accurately detected the position of anatomical layer 4. The identified alpha-beta/gamma cross-over channel was 0.08mm from the center of histologically verified layer 4 (Supplementary Fig. 4d, N=18 probes, confidence interval: -0.092mm to 0.26mm; positive values indicate distances deeper from layer 4).

**Figure 2:**
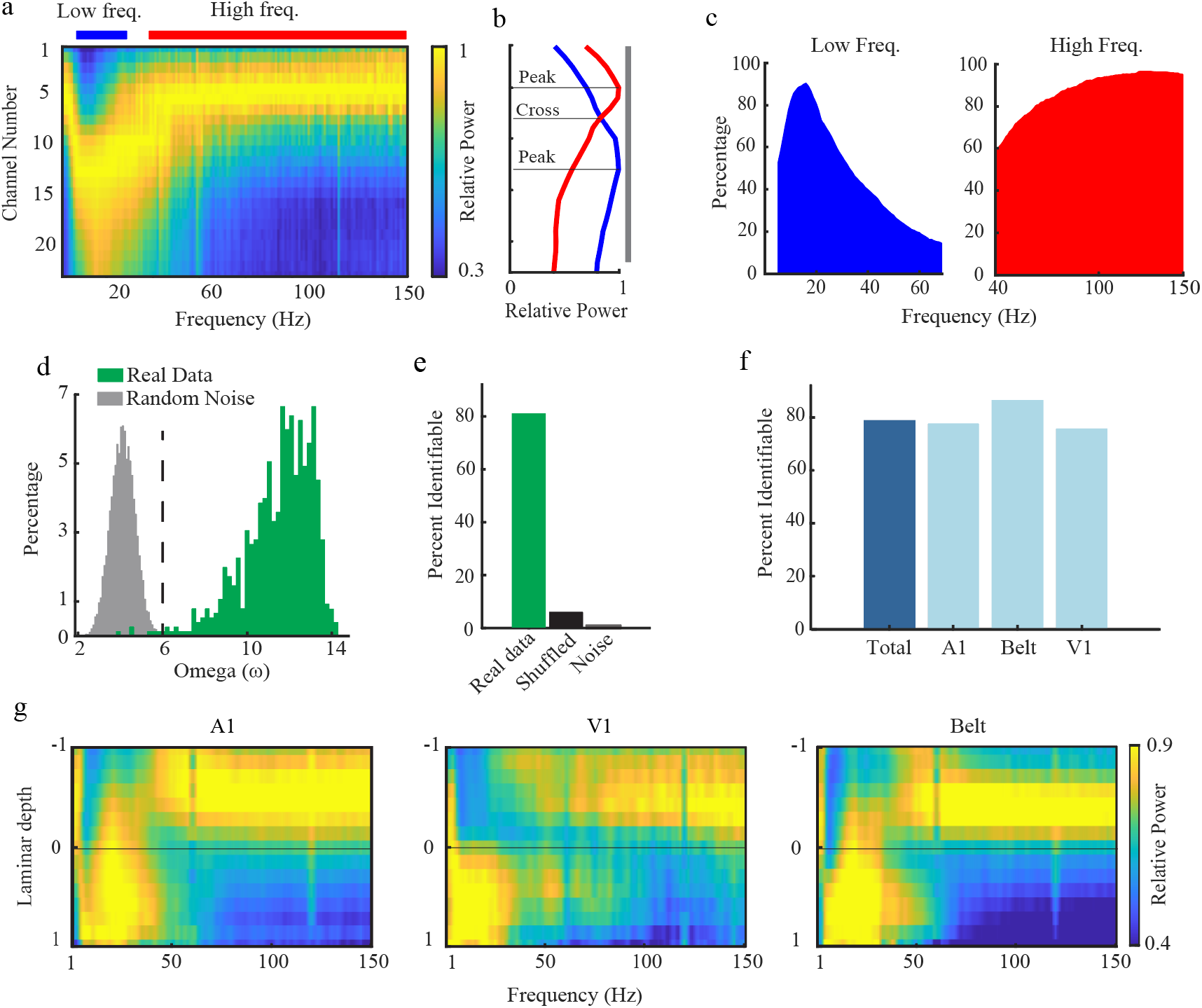
vFLIP2 algorithm validation and results. **(a**,**b)** Relative LFP power across channels and frequencies (a) and mean relative power in the low- and high-frequency ranges (b) for an example probe. The low- and high-frequency peak and crossover channels identified by vFLIP2 are marked. **(c)** percentages of probes per frequency bin that fell within the vFLIP2 low-frequency range (4–70 Hz) or high-frequency range (40–150 Hz) across all probe recordings. **(d)** Distribution of Omega (ω) among all probes in our real data and among random noise patterns. Dashed line, ω > 6 threshold. **(e)** Percentage of probes with a spectrolaminar motif identifiable by vFLIP2 in our real data, layer-shuffled data, and random noise patterns. **(f)** Percentage of probes with a spectrolaminar motif identifiable by vFLIP2 for each area in Mackey et al. **(g)** Mean spectrolaminar pattern across probes for each area in (f) as a function of laminar depth with respect to the alpha-beta/gamma crossover channel

In summary, the spectrolaminar motif is consistently identified across diverse cortical regions in data from multiple labs, including that of Mackey et al. Unlike their analysis, ours is anatomically grounded. Their feedback on our vFLIP tool helped us refine it into its next iteration, vFLIP2, which we hope will be of wide use to the community.

## Acknowledgements

We thank Jon Kaas for assistance with interpreting the histological data.

