## Supplementary Text for "A ubiquitous spectrolaminar motif across independent studies, including Mackey et al.’s own data"

This supplement provides additional responses to arguments made by Mackey et al.

Throughout the commentary, Mackey et al. criticized our use of unipolar LFP power measurements (as opposed to bipolar derivation and local referencing) and raise the topic of laminar location of alpha/beta generators/pacemakers, claiming that a method for laminar identification should “start with methods having the best available signal localization qualities.” Although we agree that the localized mechanisms of alpha/beta generators is an important topic, it is not relevant with respect to the existence nor the utility of the spectrolaminar motif. As described in our manuscript (Mendoza-Halliday et al., 2024, Extended Data Fig. 9), unipolar relative power profile provides more information, including motif orientation, than bipolar- or CSD-derived relative power.

Mackey et al. make the flawed argument that the results of our histological ground truth experiment are not reliable because “in chronic recording studies, a penetration … leaves a prominent glial scar”, which is not visible in our histological sections. Unfortunately, this argument ignores our reported methods: our recordings were acute, not chronic, and thus a “prominent glial scar” is not expected at every penetration site. Nevertheless, to provide additional documentation of our probe reconstructions, here we provide additional adjacent Nissl stained sections for an example probe (Supplementary Fig. 4) showing that the probe’s trajectory and placement can be confirmed from multiple independent sections and that the electrolytic lesions used to determine the probe’s location in cortex are clearly visible in multiple sections.

Mackey et al. also critiqued our usage of image similarity analysis. However, the ubiquity of the spectrolaminar motif does not depend on this analysis. It is present upon visual inspection of the data (Fig. 1) and can be identified using an automated algorithm (vFLIP2, Fig. 2). Our original motivation for the use of an image similarity metric was to minimize the number of assumptions about the frequencies at which we expected to find similarity. The cross-area generalization of relative power gradients in the gamma and alpha/beta ranges can also be shown statistically using Spearman rank correlation, a method free of distribution assumptions (but which requires pre-defining specific frequency bands for comparison). We previously tested whether one area’s relative power gradient in the gamma and alpha/beta predicts another area’s, amongst six frontal areas. This analysis confirmed that gamma and alpha/beta laminar gradients were positively and significantly correlated between areas, confirming ubiquity (see Supplemental Figure 2 of Bastos et al., 2018).

Mackey et al. also criticize the previous modeling work cited in our paper (Sanchez-Todo et al., 2023). The work showed that a minimal model for the spectrolaminar motif is one in which two independent generators (for gamma and alpha/beta) have distinct laminar positions. In the modeling work, the model was fit to empirical laminar data. The winning model contained a gamma generator in superficial layers and an alpha/beta generating circuit involving all layers, with an important contribution of inhibitory inputs to the apical dendrites of deep layer pyramidal neurons. We agree that further work is needed to gain more detailed knowledge of the circuits underlying the spectrolaminar motif. It is important to clarify that we cited this modeling work simply to speculate about potential mechanisms underlying the motif. Although we agree that more neurobiologically detailed models will be helpful, the model's validity is inconsequential to our discovery of the ubiquitous spectrolaminar motif.

*Additional technical description of vFLIP2*

One important advantage of the optimized computational efficiency of vFLIP2 is that it ultimately prevents a substantial incidence of false positive identifications of the spectrolaminar motif, one further concern raised by Mackey et al. This is further helped by the algorithm's use of second-degree polynomial fitting of the power gradients across the channels (filtering out statistically non-significant regressions), and the application of a threshold for our omega metric ($\omega > 6$) a more comprehensive measure of spectrolaminar pattern quality. We tested vFLIP2's false positive identification rate by analyzing the original dataset (n = 923), 1,000 channel-shuffled recordings (where the probe's laminar structure was destroyed), and 100,000 random pink noise samples like those tested by Mackey et al. vFLIP reliably identified the spectrolaminar motif in 80% of our original probe recordings while correctly rejecting identifiability in over 99% of random pink noise patterns and in over 94% of channel-shuffled probes (Figures 2d,e). In the Mackey et al. dataset, vFLIP2 identified the spectrolaminar motif in 78% of the probes (Fig. 2f). The percentage of identifiable probes was relatively consistent across areas and datasets (Supplementary Fig. 3b, Fig. 2f). For both the Mackey et al. dataset and ours, the mean spectrolaminar patterns across probes obtained with vFLIP2 (Figures 2g, Supplementary Fig. 2b,f) were remarkably similar to those obtained with the manual method (Fig. 1a-d, Supplementary Fig. 2a,e), to those aligned by the first current source density sink (Supplementary Fig. 2c,g), and in our electrolytic lesion experiment, to those aligned by the histologically-mapped layer 4 (Supplementary Fig. 2d).

Mackey et al. raised the concern that vFLIP incorrectly labeled some of their recordings as having an inverted spectrolaminar pattern. Automatic identification of inverted patterns in deep-to-superficial probe penetrations is a powerful feature of the FLIP algorithms in particular circumstances in which the orientation of the cortical sheet relative to the probe is difficult to determine (i.e., when recording from cortical folds below the cortical surface). In practice, this orientation is most often known to the electrophysiologist thanks to guidance from structural imaging and/or by comparing electrode depth to brain atlases. Taking this into consideration, vFLIP2 is now adapted to allow users to either provide the cortical orientation when known, or to let it be automatically determined when it is unknown. To test vFLIP2's automatic identification of cortical orientation, we randomly selected 100 probes with a manually-identified spectrolaminar motif, and randomly inverted the channel orientations of half of them. vFLIP2 automatically identified the correct orientation in all 100 probes.

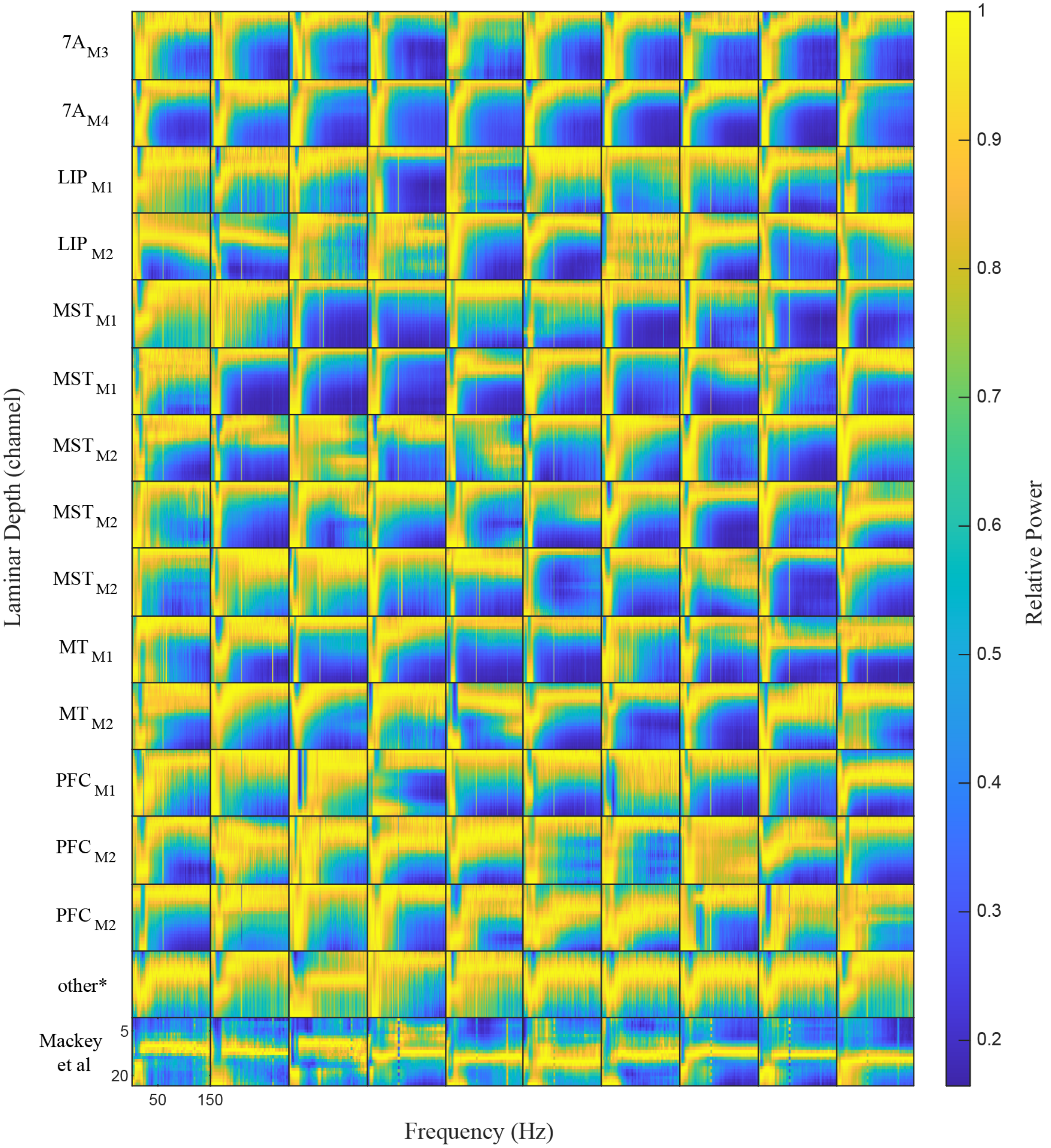

***Supplementary Figure 1****. Relative power maps as a function of frequency and laminar depth for example probes in individual areas and monkeys (subscript) from our dataset, or in Mackey et al. (including data from V1, A1, and Belt areas).*

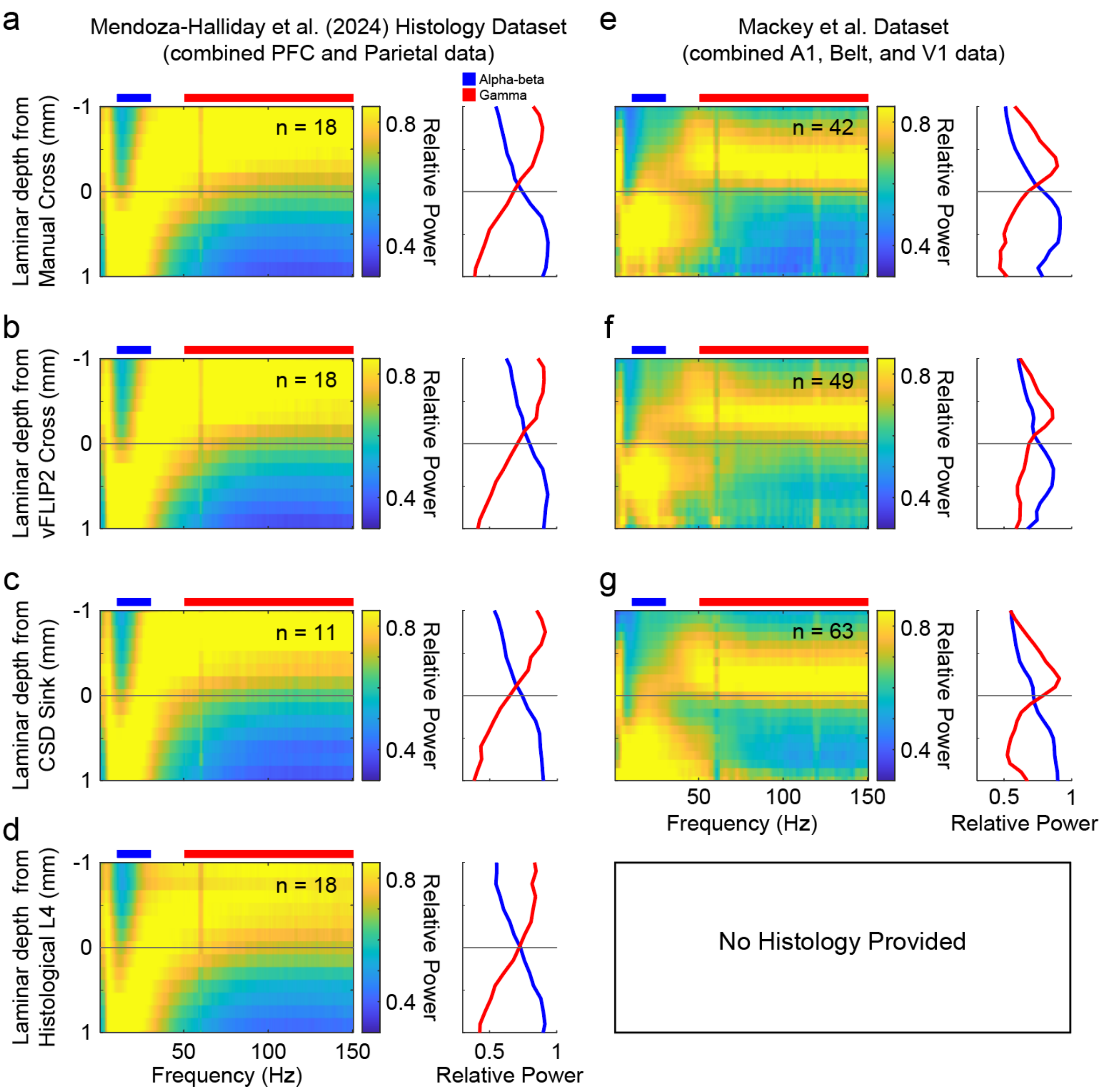


***Supplementary Figure 2. Spectrolaminar motifs are robust across different alignment techniques****. Left column, From the original publication (Mendoza-Halliday et al., 2024), data are from probes with layer 4 channels histologically verified through electrolytic lesions.* ***(a)*** *Mean relative power map and mean alpha-beta/gamma bands aligned using manual crossover*

*method.* ***(b)****, Spectrolaminar motifs are aligned using vFLIP2 algorithm.* ***(c)*** *Spectrolaminar motifs are aligned using current source density (CSD) sink.* ***(d)*** *Spectrolaminar motifs are aligned using histologically identified layer 4 channel. For each figure panel, probes were only included if alignment metric was valid, e.g., a manually identified crossover, a valid CSD profile, or an output from vFLIP2. Right column, Combined dataset from Mackey at al. dataset shows clear spectrolaminar motifs across different metrics:* ***e****, manual crossover method;* ***f****, vFLIP2 algorithm;* ***g****, CSD sink. No histological verification was provided for the Mackey et al. dataset.*

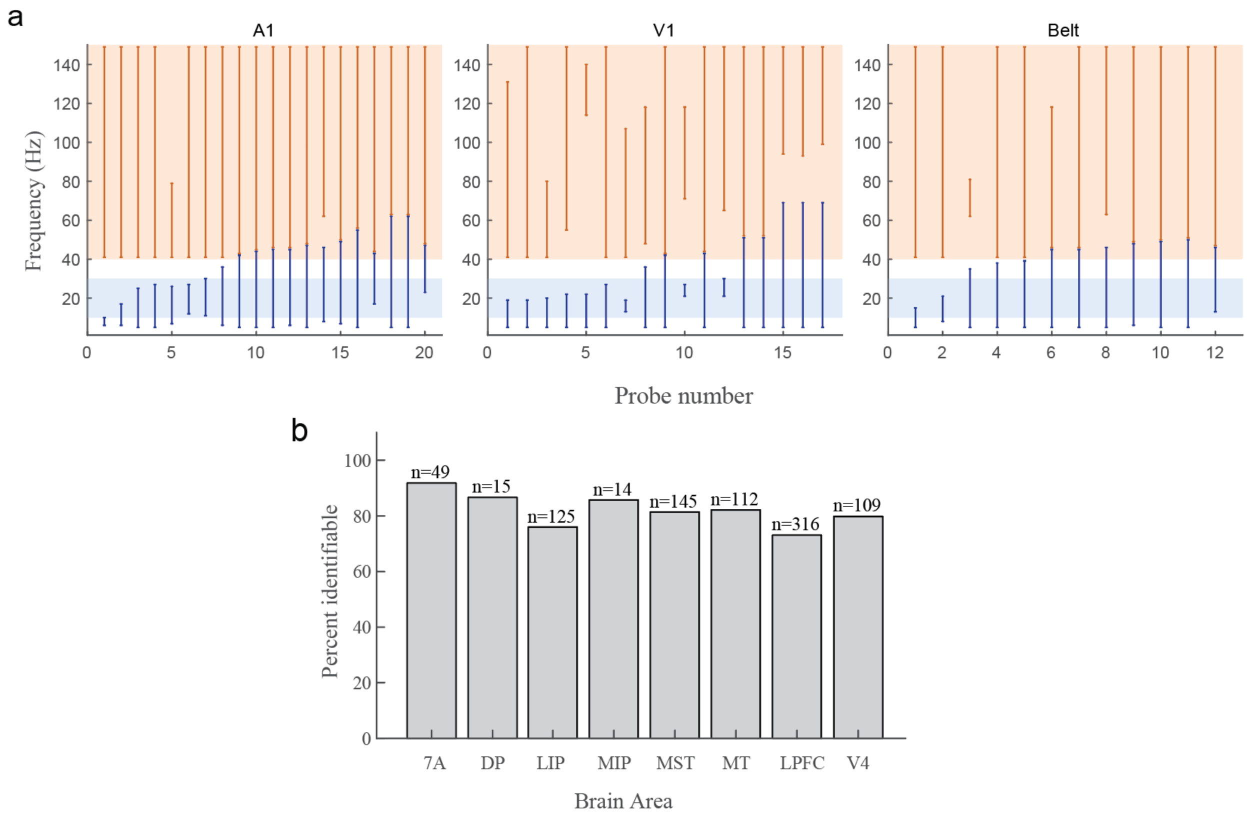


***Supplementary Figure 3. Consistency of identification of the spectrolaminar motif by vFLIP2 across probes and areas. (a)*** *Low- and high-frequency ranges showing laminar power gradients as identified by vFLIP2 in all individual probes from Mackey et al. For reference, the alpha-beta (8–30 Hz) and gamma (30-150 Hz) bands are shaded in blue and orange, respectively.* ***(b)*** *Percentage of probes with a spectrolaminar motif identifiable by vFLIP2 for each area in our data.*

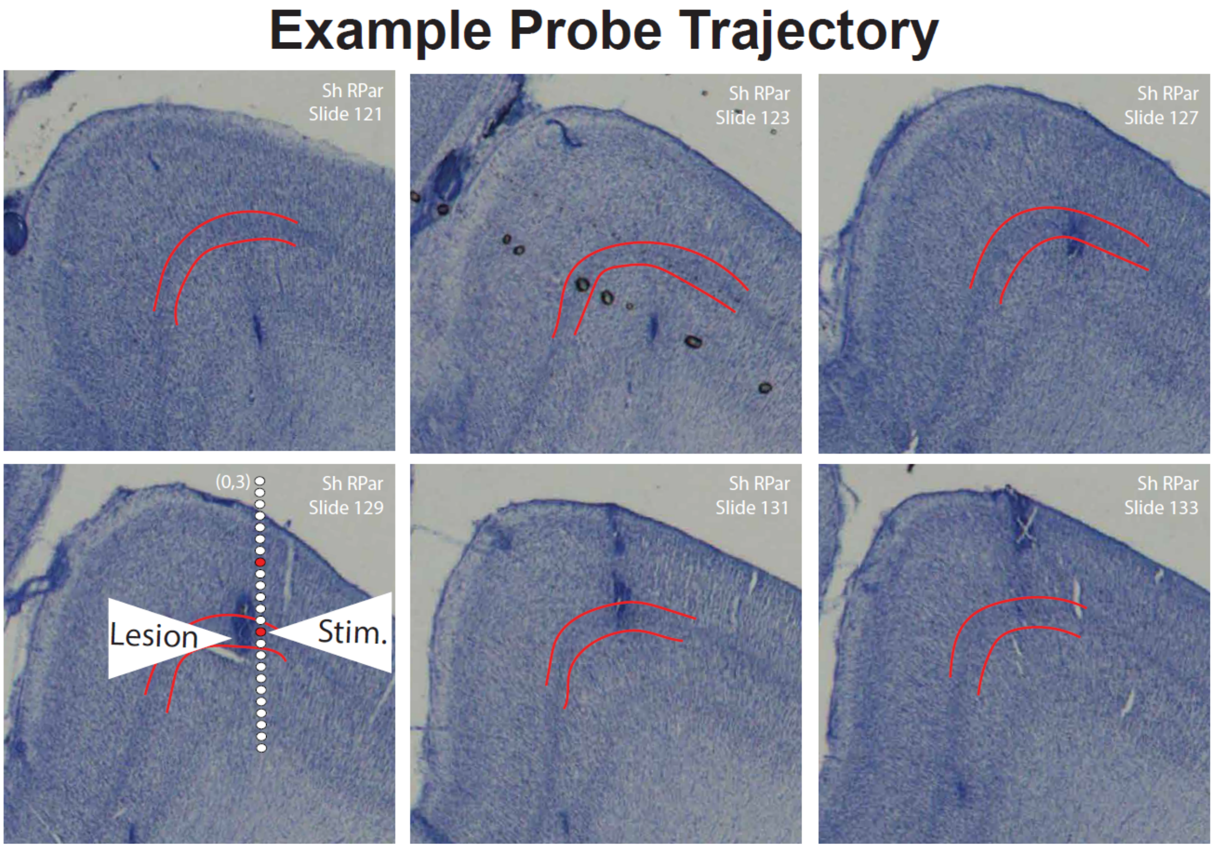


***Supplementary Figure 4. Additional histological verification of lesion locations presented in the original paper.*** *A set of adjacent histological Nissl sections from a probe in parietal cortex providing further documentation that the probe shown in our original publication in Figure 4A and Extended Data Fig. 3 was correctly reconstructed. A zoomed in version of the bottom left image was presented in the original publication. The position of the reconstructed laminar probe is shown on the bottom left (shifted slightly to the right to make the lesion visible) with the channels used for generating electrolytic lesions marked with red circles. The position of layer 4 is highlighted in red lines. Note that the position of the middle lesion used to mark the probe's position can be determined by the location of a large, darkened, punctate blob within layer 4. A secondary superficial lesion can also be identified. Each slide has a 40um thickness. Only the displayed odd sections were Nissl stained.*

| **Cortical areas reported** | **Study** |
|---|---|
| V1, V3, V4, MT, MST, 7A, DP, LIP, LPFC, MIP, Premotor Area 6, Somatosensory Area 5, Tpt, TPO | (Mendoza-Halliday et al., 2024) |
| V1, Belt, A1 | (Mackey et al., 2024) |
| VLPFC, DLPFC, 8A, ACC, SMA, 8B, DLPFC | (Bastos et al., 2018) |
| Anterior PFC, Posterior PFC | (Phillips et al., 2024), see Fig. 1f |
| V1 | (Maier et al., 2010; Smith et al., 2013; van Kerkoerle et al., 2014) |
| V1 (for superficial-layer gamma) | (Xing et al., 2012) |
| V2, TEO | (Lichtenfeld et al., 2024) |
| 8Ad, LIP | (Johnston et al., 2019) |
| SEF (for superficial-layer gamma) | (Godlove et al., 2014) |
| LIP | (Selvanayagam et al., 2024) |
| Dorsal and ventral banks of the cingulate sulcus | (Corrigan et al., 2024) |
| LPFC, MT, V4 | (Davis et al., 2023) |
| LPFC | Unpublished data from Tian Wang, Eric Kenji Lee, and Chandramouli Chandrasekaran (see Fig. 1e) |

**Supplementary Table 1:** Additional documentation of the spectrolaminar motif and its ubiquity across cortex from our original study (Mendoza-Halliday et al., 2024) as well as additional examples from the current non-human primate literature. The inclusion of these additional observations expands the ubiquity of the spectrolaminar motif from the 14 cortical areas reported in Mendoza-Halliday et al., 2024, to over 20 areas.
